# Tryptophan fluorescence quenching in β-lactam-interacting proteins is modulated by the structure of intermediates and final products of the acylation reaction

**DOI:** 10.1101/472381

**Authors:** Sébastien Triboulet, Zainab Edoo, Fabrice Compain, Clément Ourghanlian, Adrian Dupuis, Vincent Dubée, Laetitia Sutterlin, Heiner Atze, Mélanie Ethève-Quelquejeu, Jean-Emmanuel Hugonnet, Michel Arthur

**Author notes:** Corresponding Author Information. Correspondence should be addressed to and/or. Equal contribution (Sébastien Triboulet, Zainab Edoo). Equal contribution and corresponding authors (Jean-Emmanuel Hugonnet, Michel Arthur).

## Abstract

In most bacteria, β-lactam antibiotics inhibit the last cross-linking step of peptidoglycan synthesis by acylation of the active-site Ser of D,D-transpeptidases belonging to the penicillin-binding protein (PBP) family. In mycobacteria, cross-linking is mainly ensured by L,D-transpeptidases (LDTs), which are promising targets for the development of β-lactam-based therapies for multidrug-resistant tuberculosis. For this purpose, fluorescence spectroscopy is used to investigate the efficacy of LDT inactivation by β-lactams but the basis for fluorescence quenching during enzyme acylation remains unknown. In contrast to what has been reported for PBPs, we show here using a model L,D-transpeptidase (Ldt_fm_) that fluorescence quenching of Trp residues does not depend upon direct hydrophobic interaction between Trp residues and β-lactams. Rather, Trp fluorescence was quenched by the drug covalently bound to the active-site Cys residue of Ldt_fm_. Fluorescence quenching was not quantitatively determined by the size of the drug and was not specific of the thioester link connecting the β-lactam carbonyl to the catalytic Cys as quenching was also observed for acylation of the active-site Ser of β-lactamase BlaC from *M. tuberculosis*. Fluorescence quenching was extensive for reaction intermediates containing an amine anion and for acylenzymes containing an imine stabilized by mesomeric effect, but not for acylenzymes containing a protonated β-lactam nitrogen. Together, these results indicate that the extent of fluorescence quenching is determined by the status of the β-lactam nitrogen. Thus, fluorescence kinetics can provide information not only on the efficacy of enzyme inactivation but also on the structure of the covalent adducts responsible for enzyme inactivation.

---

Peptidoglycan is an essential constituent of bacterial cell walls since it prevents cell swelling and lysis by mechanically sustaining the osmotic pressure of the cytoplasm ^1^. This protective function depends upon synthesis and maintenance during the entire cell cycle of the net-like peptidoglycan macromolecule, which completely surrounds the bacterial cell. Peptidoglycan is made of glycan strands cross-linked by short peptide stems. In most bacteria, the cross-linking step is performed by D,D-transpeptidases, which are the essential targets of β-lactam antibiotics and are often referred to as penicillin-binding proteins (PBPs) ^2^. In mycobacteria ^3–4^ and in *Clostridium difficile* ^5–6^, the cross-links found are mainly (70% to 80%) formed by a second class of enzymes, the L,D-transpeptidases (LDTs). Since LDTs are not inhibited by β-lactams belonging to the penam class, such as ampicillin, ^7–8^, these enzymes are responsible for high-level resistance to these drugs in mutants of *Enterococcus faecium* ^9–10^ and *Escherichia coli* ^11^ selected in laboratory conditions.

PBPs and LDTs are structurally unrelated ^12–15^ and proceed through different catalytic mechanisms for activation of Ser and Cys nucleophiles ^16^, which are part of Lys-Ser ^17^ and Cys-His-Asp ^18^ catalytic diad and triad, respectively. PBPs and LDTs also differ by the structure of the stem peptide used as an acyl donor, a pentapeptide for PBPs ^17^ and a tetrapeptide for LDTs ^16^, except for *Enterococcus faecalis* Ldt_fs_ ^19^. PBPs cleave the D-Ala^4^-D-Ala^5^ peptide bond of the acyl donor, hence the D,D designation for these transpeptidases, and link the carbonyl of D-Ala^4^ to the side-chain amino group of the 3^rd^ residue thereby generating 4→3 cross-links ^17^ (Supplementary Fig. S1). In contrast, LDTs cleave the L-Lys^3^-D-Ala^4^ bond of the donor (L,D designation) and form 3→3 cross-links ^16^.

The first substrate of the transpeptidation reaction, the acyl donor, forms an acylenzyme intermediate that subsequently reacts with the second substrate, the acceptor, leading to the final cross-linked product. For PBPs, nucleophilic attack of the carbonyl of D-Ala^4^ by the active-site Ser results in an acylenzyme containing an ester bond and release of D-Ala^5^ ^17^. For the acylenzyme formed by LDTs, a thioester bond connects the carbonyl of L-Lys^3^ to the γ sulfur of the active-site Cys ^16^. β-lactams are structure analogues of the D-Ala^4^-D-Ala^5^ extremity of stem pentapeptides and act as suicide substrates of the PBPs ^20^. Nucleophilic attack of the carbonyl of the β-lactam ring by the active-site Ser of PBPs leads to formation of an acylenzyme, which is only hydrolyzed very slowly, leading in practice to “irreversible” inactivation in the time scale of a bacterial generation (Supplementary Fig. S1) ^2^. LDTs are also acylated by β-lactams although efficacious enzyme inactivation and antibacterial activity occur only for a single class of β-lactams, the carbapenems, such as imipenem ^7–8^. β-lactams of the cephem (cephalosporin) class, such as ceftriaxone, are only active at high concentrations since acylation is slower and the resulting thioester bond is prone to hydrolysis ^7^. The kinetic parameters are even less favorable for penams (ampicillin and penicillin) leading to equilibrium between the acylated and unacylated (functional) forms, which accounts for the lack of antibacterial activity ^7^.

Fluorescence kinetics revealed that the acylation reaction of LDTs by β-lactams of the carbapenem class comprises two limiting steps (Fig. 1A and B). In the first step, nucleophilic attack of the β-lactam carbonyl by the negatively charged sulfur atom of the catalytic Cys was thought to lead to formation of a covalent sulfur-carbon bond and to the development of a negative charge on the drug ^21^. Initially, this negative charge was proposed to be located on the β-lactam oxygen ^21^. This hypothesis was based on analogies with PBPs, which contain an oxyanion hole for stabilization of the negative charge developing on the β-lactam oxygen ^17^. However, hybrid potential simulation of the acylation reaction indicated that the path to an oxyanion is energetically disfavored in the case of LDTs and that an oxyanion could not correspond to a stabilized reaction intermediate ^22^. Rather, hybrid potential simulation indicated that the most energetically favored reaction path involves a concerted mechanism leading to the concomitant formation of the thioester bond, rupture of C-N β-lactam bond, and formation of an amine anion ^22^. In the second step, the final acylenzyme is formed by protonation of the amine anion. The origin of this proton remains elusive ^18, 22^. Formation of a non-covalent complex is not considered in this reaction scheme as the affinity of the Ldt_fm_ for carbapenems is very low, in the order of 60 to 80 mM ^7^.

**Figure 1.**
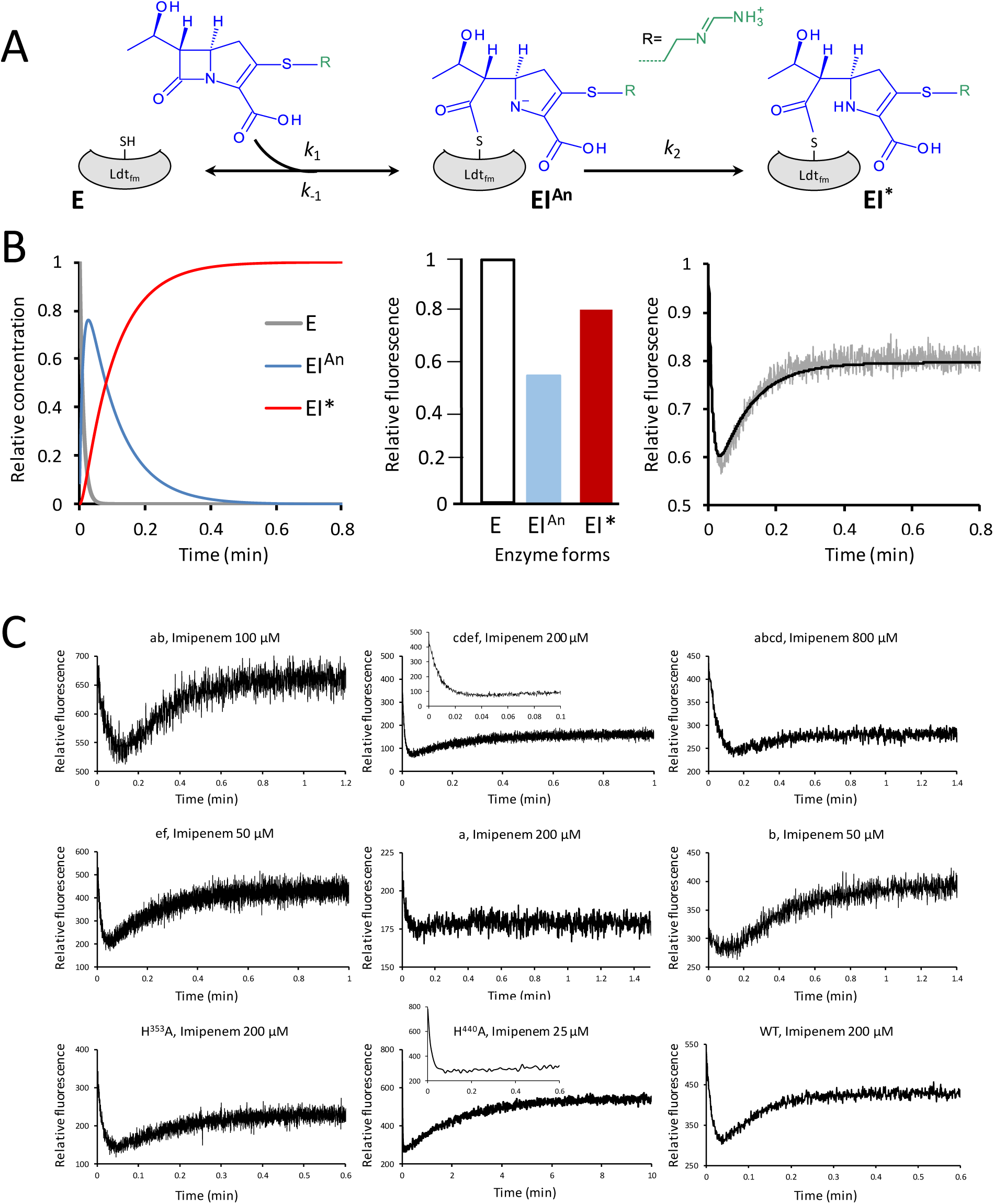
Impact of amino acid substitutions on fluorescence kinetics. (**A**) Inactivation reaction scheme. E, free Ldt_fm_; EI^An^, amine anion; EI*, acylenzyme; I, imipenem. (**B**) Fluorescence variations during Ldt_fm_ inactivation. The charts show the variations in the relative concentrations of the three enzyme forms (left panel) and the relative fluorescence intensity of the three enzyme forms (middle panel), which account for the observed fluorescence kinetics (right panel). (**C**) Fluorescence kinetics of Ldt_fm_ inactivation by imipenem. Trp residues were replaced by Phe residues in various combinations. Designation of the remaining Trp residues: a, Trp^355^; b, Trp^385^; c, Trp^410^; d, Trp^415^; e, Trp^425^; f, Trp^434^. Insets present the initial decreases of fluorescence over a short timescale.

Fluorescence kinetics proved useful to assess the efficacy of inactivation of LDTs from *Mycobacterium tuberculosis* ^23–25^ and contributed to the choice of the drugs that were tested in a phase II clinical trial ^26^. The assays were also useful in evaluating series of synthetic carbapenems ^27^ and non-β-lactam LDT inhibitors ^28^. However, the basis for the variation in fluorescence intensity detected during the two steps of the acylation reaction remained elusive. Initially, we developed fluorescence kinetics since determination of the crystal structure of Ldt_fm_ revealed a Trp residue at the entrance of the catalytic cavity ^15^, which appeared ideally located for fluorescence quenching upon drug binding due to changes in its environment ^21^. However, this naïve explanation was subsequently found to be insufficient to account for the various behaviors observed for acylation of LDTs by representatives of the β-lactam classes ^7, 22-25^. This prompted us to investigate here the basis for variations in fluorescence intensity occurring upon acylation of LDTs using the well-characterized L,D-transpeptidase Ldt_fm_ from *E. faecium* as a model. We show that variations in fluorescence intensity are not due to modification of the environment of Trp residues but to the formation of a β-lactam-derived fluorescence quencher during the acylation reaction.

## RESULTS

### Trp to Phe and His to Ala substitutions do not abolish the biphasic behavior of fluorescence kinetics observed for Ldt_fm_ inactivation by imipenem

We have previously proposed that the acylation of Ldt_fm_ by carbapenems is a two-step reaction involving reversible formation of an amine anion (EI^An^, step 1) followed by irreversible formation of an acylenzyme (EI*, step 2) (Fig. 1A) ^21–22^. Fluorescence kinetics also display two phases enabling the determination of kinetic parameters for the two steps of the reaction (Fig. 1B) ^21^. In the first phase, fluorescence intensity decreases because EI^An^ is rapidly formed to the detriment of the free enzyme (E) and the fluorescence intensity of EI^An^ is lower than that of E. In the second phase, fluorescence intensity increases since the acylenzyme (EI*) is formed to the detriment of EI^An^ (acylation step) and the fluorescence intensity of EI* is greater than that of EI^An^. Our first aim was to determine whether variations in the fluorescence intensity could be correlated to the modification of the environment of a specific Trp residue of Ldt_fm_. To address this question, Trp residues of the catalytic domain of Ldt_fm_ were replaced by Phe residues in various combinations. For the sake of simplicity, the six Trp residues at positions 355, 385, 410, 415, 425, and 434 of Ldt_fm_ were designated as residues a to f, respectively. Fluorescence kinetics were determined for Ldt_fm_ derivatives lacking 2 to 5 Trp residues (Fig. 1C). Fluorescence kinetics remained bi-phasic in all cases, except for the Ldt_fm_ derivative only retaining the “a” residue (*i*.*e*. Trp355). Thus, the variations in the fluorescence intensity associated with the two steps of the acylation reaction do not depend upon the modification of the environment of a specific Trp residue of Ldt_fm_. This prompted us to investigate fluorescence quenching by His residues of Ldt_fm_. Fluorescence kinetics obtained with the His^353^Ala and His^440^Ala derivatives of Ldt_fm_ indicated that neither residue is essential for biphasic fluorescence kinetics (Fig. 1C). His at position 421 could not be analyzed since this residue is part of the catalytic triad and is essential for Ldt_fm_ activity.

### Impact of Trp to Phe and His to Ala substitutions on the efficacy of the inactivation reaction and on the relative fluorescence intensity of the three enzyme forms

Fitting simulations to experimental data for four concentrations of imipenem was used to determine the inactivation kinetic parameters (Supplementary Fig. S2). None of the Trp residues was essential for the acylation of Ldt_fm_ by imipenem although important variations were observed in the *k*_1_ (formation of the amine anion) and *k*_2_ (acylation step) parameters (Fig. 2A and 2B, respectively). The Trp to Phe substitutions produced large variations in the intrinsic fluorescence of the three forms of the enzyme (apo Ldt_fm_, amine anion, and acylenzyme; Fig. 2C). The formation of the acylenzyme (EI*) was confirmed by mass spectrometry for all Ldt_fm_ derivatives (mass increment of 299.3 Da; data not shown).

**Figure 2.**
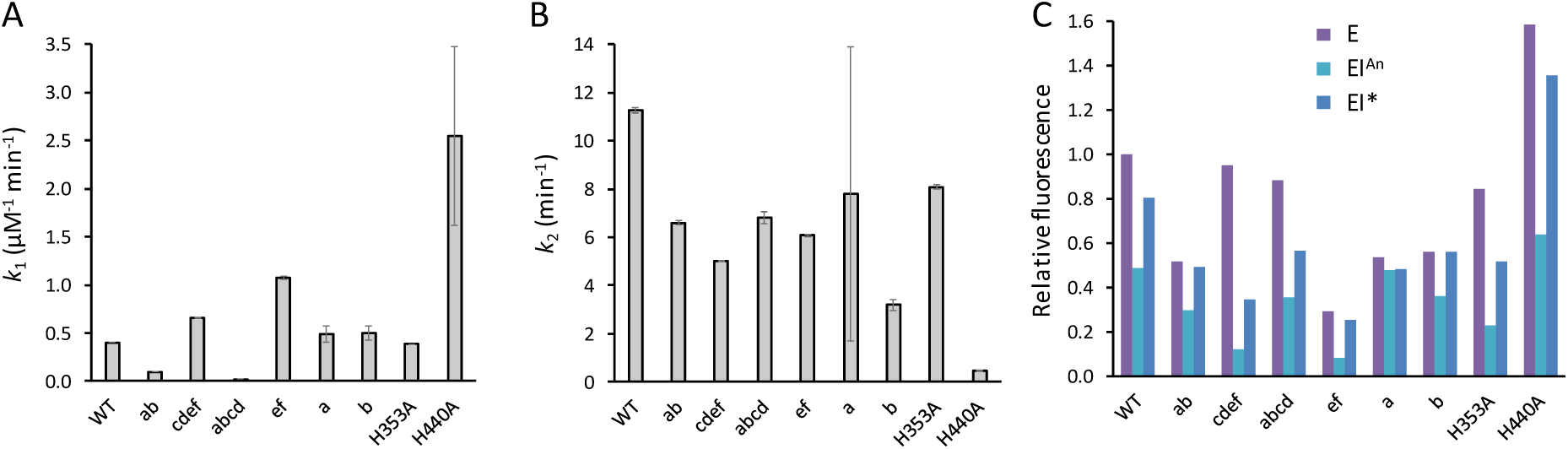
Inactivation efficacy and relative fluorescence intensity of Ldt_fm_ and derivatives with Trp to Phe and His to Ala substitutions. (**A**) Kinetic parameter *k*_1_ for formation of the amine anion. (**B**) Kinetic parameter *k*_2_ for the acylation step. Error bars represent standard deviations from the fitted curves. (**C**) Fluorescence intensity of the three enzyme forms relative to that of the free form of the wild-type enzyme. E, Ldt_fm_; EI^An^, amine anion; EI*, acylenzyme. All, presence of the full complement of Trp residues (a, d, c, d, e, and f). Trp residues were replaced by Phe residues in various combinations. Designation of the remaining Trp residues: a, Trp^355^; b, Trp^385^; c, Trp^410^; d, Trp^415^; e, Trp^425^; f, Trp^434^.

### Conservation of Trp residues in Ldt_fm_ homologues

The Trp residues were poorly conserved in L,D-transpeptidases from Gram-positive and Gram-negative bacteria (Supplementary Fig. S3), as expected from the substantial high residual acylation activity of Ldt_fm_ derivatives containing Trp to Phe substitutions (above, Fig. 2A and B). However, residue e (Trp^425^) was relatively conserved in L,D-transpeptidases from *M. tuberculosis*, being present in the Mt1, Mt2, Mt3, and Mt4 enzymes as well as in the corresponding orthologues of *M. abscessus*. These enzymes are known to display bi-phasic fluorescence kinetics upon inactivation by carbapenems ^23–24, 29^ and (unpublished results). Since Trp^425^ was the only conserved Trp residue in the LDTs from mycobacteria we investigated the impact of a Trp to Phe substitution at this position in Ldt_fm_. As shown in Supplementary Fig. S4A and B, derivatives of Ldt_fm_ lacking all Trp residues except Trp^425^ or lacking only this residue displayed biphasic kinetics. These results indicated that the biphasic nature of fluorescence kinetics does not depend upon any conserved Trp residue in the L,D-transpeptidase protein family.

### Fluorescence kinetics with a single ectopic Trp residue at position 383

All six Trp residues of Ldt_fm_ were replaced by Phe residues. As expected, the variant devoid of Trp residues was very weakly fluorescent (data not shown). No modification of this weak fluorescence was observed upon acylation by imipenem. This observation ruled out the formal possibility that the catalytic Cys acylated by imipenem could be a fluorophore. Replacement of Tyr at position 383 by a Trp residue in this Ldt_fm_ derivative restored biphasic fluorescence kinetics (Supplementary Fig. S4C). This result confirmed that biphasic fluorescence kinetics are not dependent upon the presence of any of the six Trp residues of Ldt_fm_ in their original positions.

### Investigation of the opened β-lactam ring linked to the catalytic Cys residue as a quencher

Since the variations in the fluorescence observed during the two steps of the acylation reaction could not be accounted for by modification of the environment of any specific Trp residue, our next objective was to investigate whether quenching could result from energy transfer from Trp residues to the Cys-β-lactam covalent adduct generated upon enzyme inactivation. To investigate this possibility, we first examined acylation of Ldt_fm_ by the chromogenic cephalosporin nitrocefin. Fluorescence kinetics with this compound displayed a monophasic behavior with a large decrease (72%) in the fluorescence intensity (Fig. 3A). In the Ldt_fm_-nitrocefin acylenzyme, the negative charge developing on the β-lactam nitrogen is stabilized by a mesomeric effect that prevents protonation of the amine ^30^. Thus, the amine anion formed as an end product with nitrocefin (Fig. 3A) or as an intermediate with imipenem (Fig. 1) was associated with extensive fluorescence quenching. As found for nitrocefin, acylation of Ldt_fm_ by ceftriaxone led to a monophasic decrease in the fluorescence intensity (Fig. 3B). Acylation of Ldt_fm_ by ceftriaxone involves a concerted mechanism leading to the elimination of the side chain and formation of a conjugated imine ^7^. The extent of fluorescence quenching observed for the acylenzyme formed with ceftriaxone (40%; imine) or by the intermediate formed with imipenem (45%; amine anion) were more important than that observed for the final Ldt_fm_-imipenem adduct (20%) suggesting that quenching was decreased by protonation of the β-lactam nitrogen.

**Figure 3.**
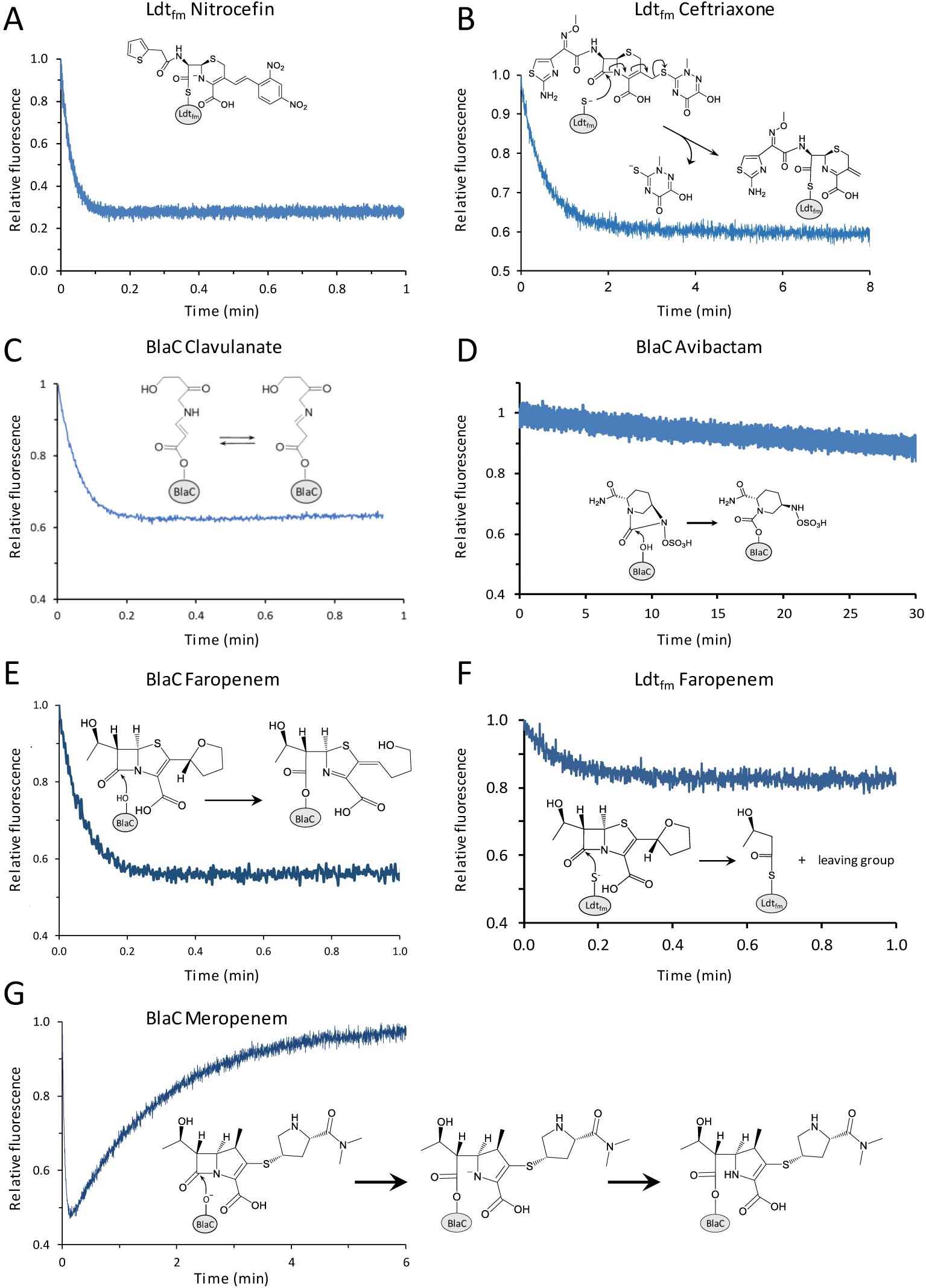
Variations in fluorescence intensity resulting from acylation of L,D-transpeptidase Ldt_fm_ and β-lactamase BlaC by various β-lactams and β-lactamase inhibitors. Stopped-flow fluorescence kinetics were obtained with final concentrations of 10 µM for the enzymes and 100 µM for β-lactams and β-lactamase inhibitors (**A**) The negative charge developing on the amine nitrogen upon acylation of Ldt_fm_ by nitrocefin is stabilized by a mesomeric effect involving the electron withdrawing dinitrobenzene ^30, 42^. (**B**) Transfer of the negative charge to a leaving group in the concerted mechanism leading to acylation of Ldt_fm_ by ceftriaxone ^7^. (**C**) Tautomeric forms of the acylenzyme resulting from acylation of BlaC by clavulanate and decarboxylation of the β-lactamase inhibitor ^32, 41^. (**D**) BlaC-avibactam carbamoylenzyme 33. (**E** and **F**) Acylation of BlaC and Ldt_fm_ by faropenem, respectively ^24^. (**G**) Accumulation of an amine anion in the onset of the reaction of hydrolysis of meropenem by BlaC (BlaC^An^). This is followed by protonation of the β-lactam nitrogen ^35^.

### Fluorescence quenching associated with inactivation of β-lactamases by β-lactamase inhibitors

In order to enrich the structural diversity of molecules investigated in the fluorescence quenching assay, we aimed to test β-lactamase inhibitors. Most β-lactams could not be tested with β-lactamases since the fluorescence assay requires relatively high enzyme concentrations (typically 10 µM) leading to rapid hydrolysis of the drugs even if the turnovers are low. For this reason, we focused on inactivation of BlaC from *Mycobacterium tuberculosis* by clavulanate since the corresponding acylenzyme is not prone to hydrolysis at the timescale of our experiments ^31^. Acylation of BlaC by clavulanate led to a large decrease in the fluorescence intensity (37%) (Fig. 3C). Previous studies have shown that acylation of BlaC by clavulanate leads to formation of imine and *trans*-enamine tautomers (depicted in Fig. 3C) following drug decarboxylation ^32^. Since Ldt_fm_ and BlaC contain Cys and Ser as the active-site nucleophile, respectively, these results indicate that fluorescence quenching is observed with acylenzymes containing both thioester and ester bonds.

Avibactam offered the possibility to test a non-β-lactam inhibitor that inhibits BlaC by formation of a carbamoylenzyme ^33^. Formation of the adduct (Supplementary Fig. S5) was not associated with any significant modification of the fluorescence intensity of the enzyme (Fig. 3D). Protonation of the amino sulfate nitrogen of avibactam in the avibactam-BlaC adduct was thus associated with limited quenching as found for the protonation of the β-lactam nitrogen in the final Ldt_fm_-imipenem and BlaC-clavulanate adducts (Fig. 1 and 3C).

### Acylation of Ldt_fm_ and BlaC by faropenem

Faropenem, a β-lactam belonging to the penem class, is poorly hydrolyzed by BlaC and rapidly inactivates L,D-transpeptidases ^24^. Thus, faropenem provided the opportunity to study the acylation of the two enzyme types, *i*.*e*. a β-lactamase (BlaC) and an L,D-transpeptidase (Ldt_fm_), by the same β-lactam. Acylation of BlaC by faropenem led to extensive fluorescence quenching (44%) (Fig. 3E). The resulting acylenzyme contained an unprotonated β-lactam nitrogen. In contrast, acylation of Ldt_fm_ by faropenem, which resulted in the rupture of the C^5^-C^6^ bond and the loss of a large portion of the drug including the β-lactam nitrogen ^24^, was associated with a moderate decrease in the fluorescence intensity (17%) (Fig. 3F).

### Acylation of BlaC by the meropenem carbapenem

Although BlaC displays moderate carbapenemase activity, previous studies have shown that a BlaC-meropenem adduct is the predominant enzyme form upon incubation of the enzyme with low drug concentrations ^34–35^. As expected, the control experiment depicted in Supplementary Fig. S6 shows that the concentration of meropenem remains saturating for at least 80 min during hydrolysis of meropenem (40 – 200 µM) by BlaC (10 µM). Fluorescence kinetics at a shorter time scale (6 min) showed a transitory decrease in fluorescence intensity (40%) (Fig. 3G). This observation suggests that formation of an amine anion initially leads to an important fluorescence quenching. Then, fluorescence returned to the initial level suggesting that protonation of the amine anion fully suppressed quenching. Thus, formation of an amine anion following nucleophilic attack of the carbonyl carbon of carbapenems by Ldt_fm_ (Fig. 1) and BlaC (Fig. 3G) similarly led to extensive fluorescence quenching despite the presence of different catalytic residues (Cys *versus* Ser) and different modes of activation of these nucleophiles involving His and Lys residues.

## DISCUSSION

The main aim of the present study was to determine the basis for the variations in the fluorescence intensity observed upon acylation of LDTs by β-lactam antibiotics. Previous analyses of the D,D-transpeptidase R61 (PBP family) have shown that acylation of the enzyme by penicillin G leads to a fluorescence quenching, which specifically depends upon one of two Trp residues of the protein ^36–37^. This residue, W^233^, is essential for the D,D-carboxypeptidase activity of the protein, as determined by the release of the terminal D-Ala residue from a model peptide ending in D-Ala-D-Ala ^37^. The crystal structure of R61 in complex with a peptide mimicking the acyl donor revealed hydrophobic interactions between the side-chain of W^233^ and methylene groups of this substrate ^38^. Likewise, the structure of a penicillin-R61 complex revealed that W^233^ is in hydrophobic interaction with the phenylacetyl side chain of the drug ^39^ indicating that fluorescence quenching may depend upon this direct interaction. In contrast, we show here that the variations in fluorescence during acylation of LDTs by β-lactam antibiotics require at least one Trp residue but that the position of this residue is not determinant. This was established by deleting Trp residues from Ldt_fm_ in various combinations (Fig. 1 and 2), by introducing a single Trp residue in an ectopic position of Ldt_fm_ (Fig. S4), and by showing that large variations in fluorescence intensity occur in distantly related LDTs and unrelated β-lactamases that do not share any conserved Trp residue (Fig. S3). Since fluorescence quenching was not mainly due to changes in the environment of Trp residues, we investigated the possibility that the nucleophilic attack of the β-lactam ring could itself generate a fluorescence quencher. If this is the case, the fluorescence profile should depend upon the structural features of the drug bound to the active-site residue. Accordingly, extensive quenching was not mainly determined by the size of the side chains of β-lactams or by the nature of the enzyme nucleophile (Cys *versus* Ser in Ldt_fm_ and BlaC, respectively) (Fig. 3). Acylation of Ser or Cys *per se* was not sufficient for fluorescence quenching since a modest or no decrease in fluorescence intensity was observed for the final acylation products of Ldt_fm_ by faropenem, of BlaC by avibactam, and of both enzymes by carbapenems. By elimination, these results pointed to the status of the β-lactam nitrogen as the only source of variation that could account for the main variations in fluorescence intensity. A large fluorescence quenching occurred if the β-lactam nitrogen was negatively charged, as transiently found during acylation of Ldt_fm_ and BlaC by carbapenems. A large fluorescence quenching also occurred if the nitrogen atom was engaged in a double bond as observed for the acylenzymes formed by Ldt_fm_ with ceftriaxone or nitrocefin and by BlaC with faropenem or clavulanate. In agreement with the critical role of the double bond, fluorescence quenching was not observed for acylation of BlaC by avibactam, which results in the formation of a carbamoyl-enzyme with a protonated nitrogen, or for acylation of Ldt_fm_ by faropenem, which results in the formation of an acylenzyme lacking the nitrogen atom following rupture of the C^5^-C^6^ bond. These results indicate that fluorescence kinetics can provide information not only on the efficacy of enzyme inactivation but also on the structure of the covalent adducts, including for example the presence of Δ1- or Δ2-pyrroline ring in enzyme-carbapenem adducts. Our results also indicate that nitrocefin provides a very sensitive assay to titrate the active site of enzymes that do not hydrolyze this cephalosporin. This property could be exploited to identify β-lactams and non-β-lactam inhibitors that reversibly or irreversibly bind to LDTs.

## EXPERIMENTAL SECTION

### Enzyme production and purification

The catalytic domain of Ldt_fm_ (residues 341 to 466) and soluble BlaC (residues 39 to 306) were produced in *E. coli* and purified by metal affinity and size exclusion chromatography as previously described ^30^. Synthetic genes were purchased for production of Ldt_fm_ derivatives with Trp to Phe and His to Ala substitutions (GeneCust). All experiments were performed with a fixed concentration of Ldt_fm_ and BlaC (10 µM).

### Spectrofluorimetry

Fluorescence kinetics were studied in 100 mM sodium phosphate (pH 6.0) at 20°C by using a stopped-flow apparatus (RX-2000, Applied Biophysics) coupled to a spectrofluorimeter (Cary Eclipse; Varian). Excitation was performed at 224 nm with a 5 nm slit and a 2 mm optical path. Fluorescence emission was determined at 335 nm with a 5 nm slit and a 10 mm optical path. Fluorescence data were collected exactly in the same conditions for comparison of the relative fluorescence of wild-type Ldt_fm_ and derivatives with Trp to Phe and His to Ala substitutions. These conditions included the photomultiplier voltage, which was set to 600 V. Kinetic constants were determined using the DynaFit software (BioKin Ltd) 40 for each Ldt_fm_ derivative by simultaneously fitting the fluorescence data for the various inhibitor concentrations to differential equations derived from the reaction scheme depicted in Fig. 1A.

### Mass spectrometry

Formation of the proposed Ldt_fm_-β-lactam and BlaC-β-lactam adducts in the conditions described in the text was systematically checked by mass spectrometry (data not shown). Enzymes and β-lactams were incubated for appropriate time periods at 20°C. Five μl of acetonitrile and 1 μl of 1% formic acid were extemporaneously added. Injection in the mass spectrometer (Qstar Pulsar I; Applied Biosystem) was performed at a flow rate of 0.05 ml/min (acetonitrile, 50%, water, 49.5%, and formic acid, 0.5%; per volume). Spectra were acquired in the positive mode, as previously described ^8^. Detailed analyses of the mass spectra of the β-lactam adducts considered in the current study have been previously published ^7–8, 24, 29, 41^.

## ANCILLARY INFORMATION

### Supporting Information

Supplementary figures S1 to S6.

### Author Contribution

ST, ZE contributed equally to this work. MA and JEH contributed equally to this work.

## Acknowledgement

We thank L. Dubost and A. Marie for technical assistance in the collection of mass spectra. This work was supported by the French National Research Agency (grant MycWall, ANR-17-CE18-0010) and the Fondation pour la Recherche Médicale (grant ECO20160736080 to ZE).

