## Supplementary Information for "Tryptophan fluorescence quenching in β-lactam-interacting proteins is modulated by the structure of intermediates and final products of the acylation reaction"

### SUPPORTING INFORMATION

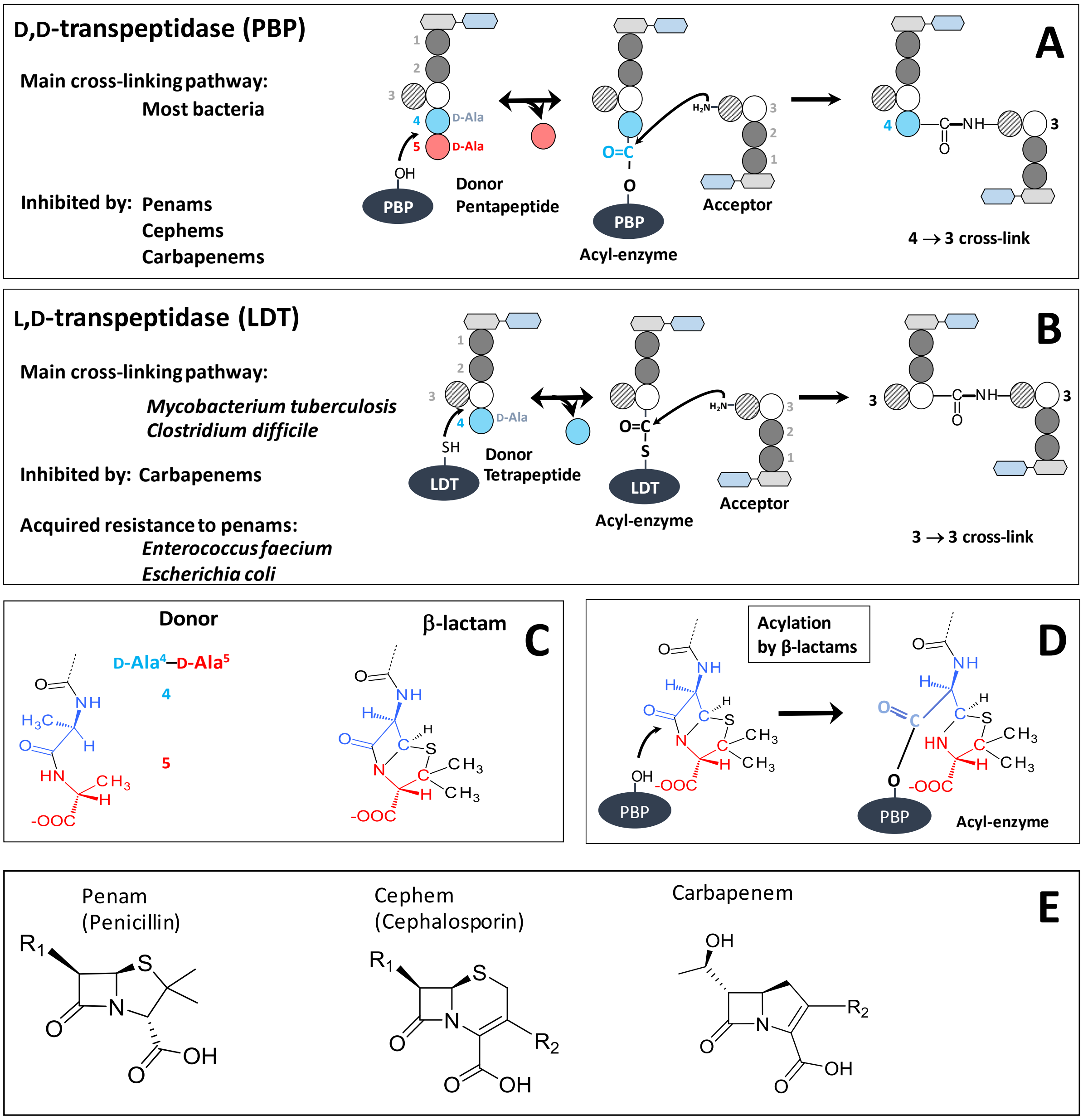


**Supplementary Fig. S1. Reactions catalyzed by PBPs and LDTs with peptidoglycan precursors and β-lactams.** (**A**) and (**B**) Peptidoglycan cross-linking by PBPs and LDTs, respectively. (**C**) Structural similarity between β-lactams and the D-Ala-D-Ala extremity of peptidoglycan precursors. (**D**) Acylation of a PBP by a β-lactam (penam). (**E**) Structure of the three main classes of β-lactams.

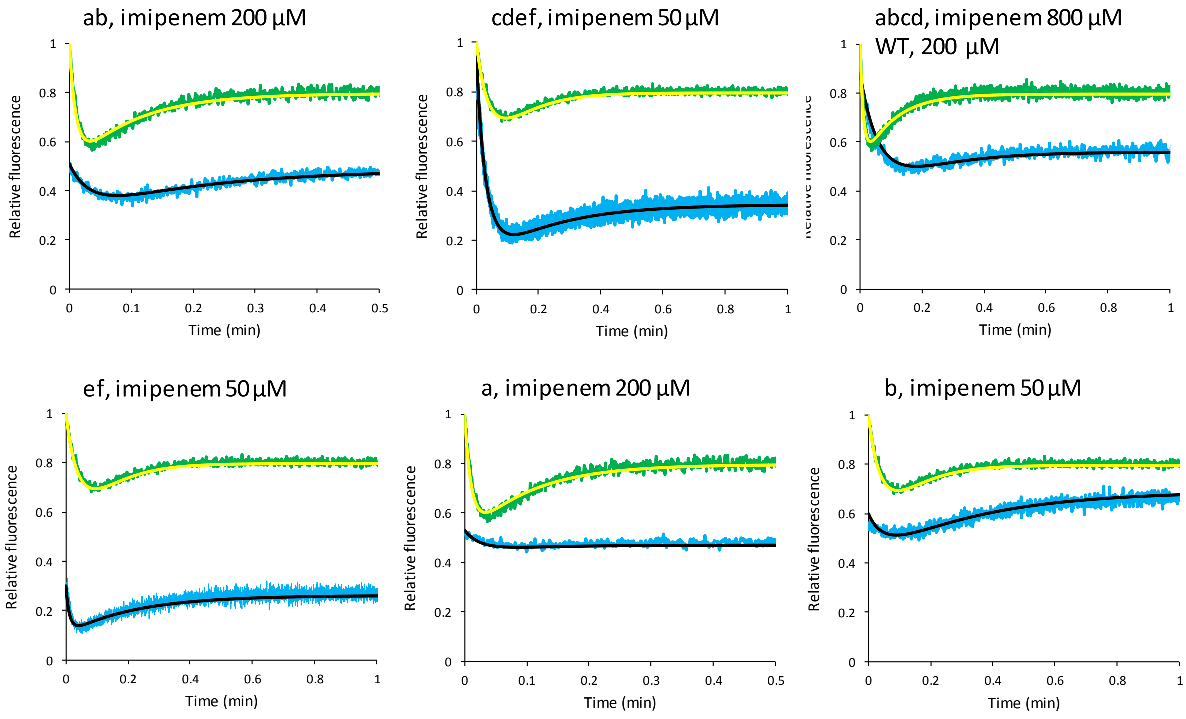


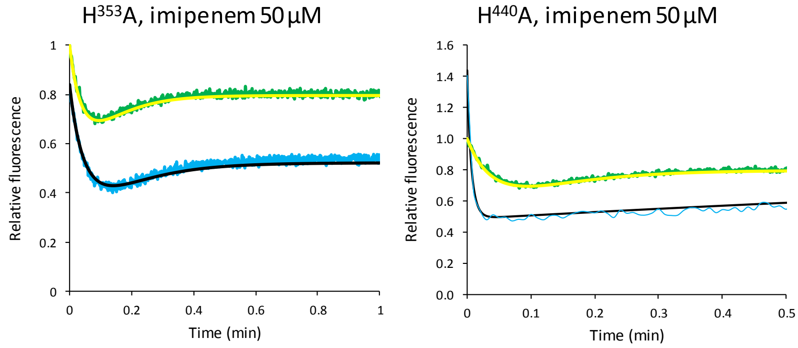


Trp designation

| Trp | position |
|---|---|
| a | 355 |
| b | 385 |
| c | 410 |
| d | 415 |
| e | 425 |
| f | 434 |

**Supplementary Fig. S2. Fluorescence kinetics for inactivation of Ldt$_{fm}$ and derivatives by imipenem.** Simulations (black line) were fitted to experimental data (blue) for eight derivatives of Ldt$_{fm}$ with various substitutions. Data and fit for the parental enzyme are indicated in each panel (green and yellow, respectively). The figure shows representative kinetics for one of the four concentrations of imipenem that were tested for each enzyme.

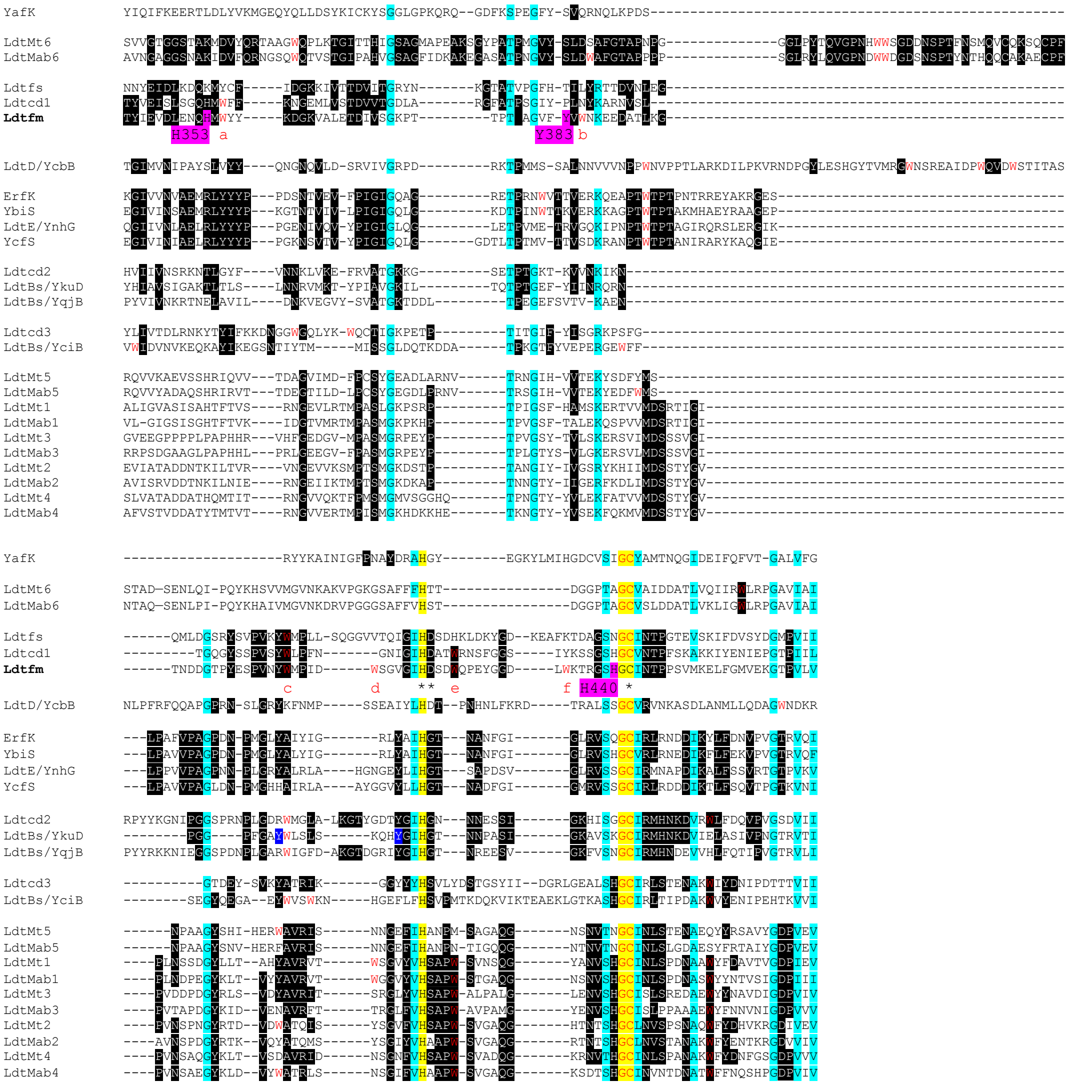


**Supplementary Fig. S3. Alignment of the amino acid sequence of the catalytic domain of various L,D-transpeptidases.** Enzymes originate from *Mycobacterium tuberculosis* ($Ldt_{Mt1}$ to $Ldt_{Mt6}$), *Mycobacterium abscessus* ($Ldt_{Mab1}$ to $Ldt_{Mab6}$), *Enterococcus faecium* ($Ldt_{fm}$), *Enterococcus faecalis* ($Ldt_{fs}$), *Escherichia coli* (YcbB, YnhG, YafK, Ybis, ErfK, and YcfS), *Bacillus subtilis* (YkuD, YqjB, and YciB) and *Clostridium difficile* ($Ldt_{Cd1}$ to $Ldt_{Cd3}$). Highlights, yellow, invariant residues; blue, highly conserved residues across all sequences; black, conserved residues in subgroups of sequences. Trp residues are indicated in red with the a, b, c, d, e, and f designation for $Ldt_{fm}$. The His and Tyr residues of $Ldt_{fm}$ that were replaced by Ala and Trp, respectively, are highlighted in pink. Stars indicate residues of the His-Asp-Cys catalytic triad.

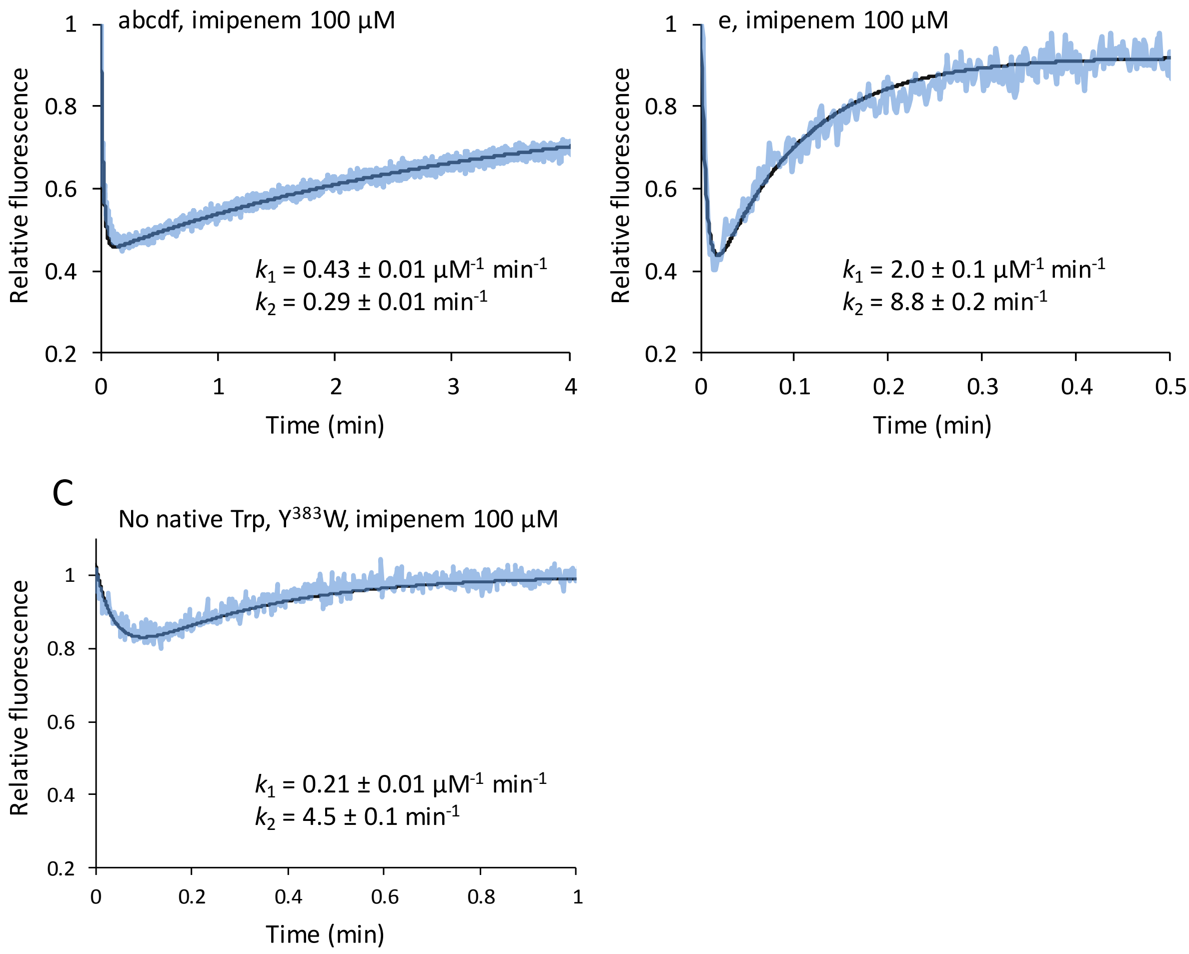


**Supplementary Fig. S4. Fluorescence kinetics of Ldt$_{fm}$ derivatives.** (**A** and **B**) Role of Trp$^{425}$ (residue e) which is conserved in L,D-transpeptidases from mycobacteria. Kinetics were performed for Ldt$_{fm}$ derivatives containing all Trp residues except Trp$^{425}$ (residues a, b, c, d, and f present) or lacking all Trp residues except Trp$^{425}$ (residue e present). (**C**) Kinetics for an Ldt$_{fm}$ derivative lacking all Trp residues in their original position and containing a single Trp residue at an ectopic position (Tyr to Trp substitution at position 383).

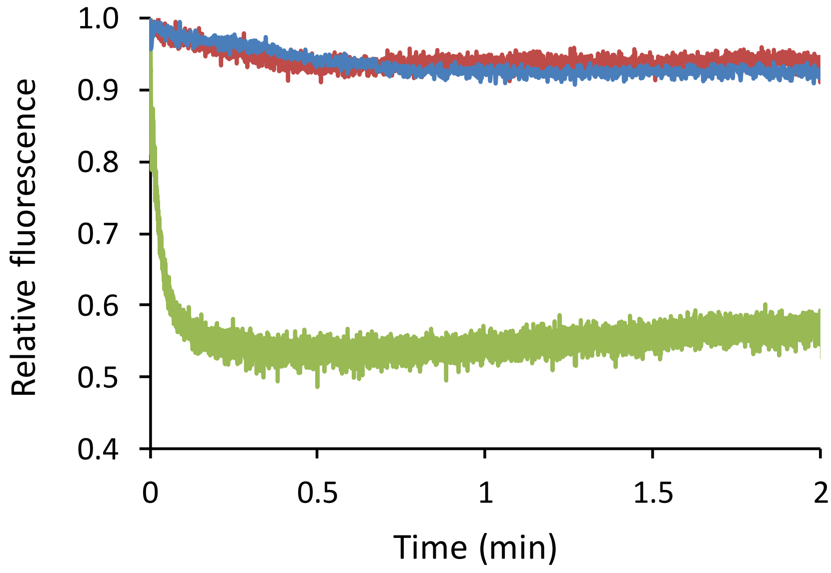


**Supplementary Fig. S5. Titration of free BlaC with clavulanate.** Green, stopped-flow fluorescence kinetics showing the fluorescence quenching due to acylation of BlaC (10 µM) by clavulanate (300 µM). Blue, pre-incubation of BlaC (10 µM) with avibactam (100 µM) fully abolished fluorescence quenching. Red, control containing only BlaC (10 µM). These data show that BlaC is fully acylated by avibactam.

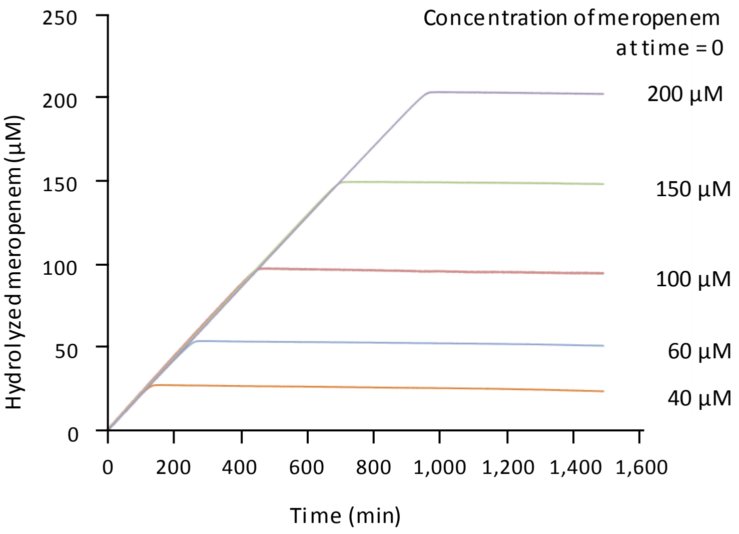


**Supplementary Figure S6. Hydrolysis of meropenem by BlaC.** This control experiment shows that BlaC (10 µM) remains saturated by meropenem (40 to 200 µM) during extended time periods (90 min for the lowest concentration). This is due to a combination of low turnover number ($k_{cat}$ = 0.08 ± 0.01 $min^{-1}$) and $K_m$ (3.4 µM ± 0.7) values (34). Hydrolysis of meropenem was monitored by spectrophotometry ($\Delta\varepsilon_{298\ nm}$ = -7,700 $M^{-1}$ $cm^{-1}$).
